# Evaluating the use of ABBA-BABA statistics to locate introgressed loci

**DOI:** 10.1101/001347

**Authors:** Simon H. Martin, John W. Davey, Chris D. Jiggins

## Abstract

Several methods have been proposed to test for introgression across genomes. One method tests for a genome-wide excess of shared derived alleles between taxa using Patterson’s *D* statistic, but does not establish which loci show such an excess or whether the excess is due to introgression or ancestral population structure. Several recent studies have extended the use of *D* by applying the statistic to small genomic regions, rather than genome-wide. Here, we use simulations and whole genome data from *Heliconius* butterflies to investigate the behavior of *D* in small genomic regions. We find that *D* is unreliable in this situation as it gives inflated values when effective population size is low, causing *D* outliers to cluster in genomic regions of reduced diversity. As an alternative, we propose a related statistic 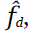 a modified version of a statistic originally developed to estimate the genome-wide fraction of admixture. 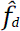 is not subject to the same biases as *D*, and is better at identifying introgressed loci. Finally, we show that both *D* and 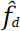 outliers tend to cluster in regions of low absolute divergence (*d_XY_*), which can confound a recently proposed test for differentiating introgression from shared ancestral variation at individual loci.

## INTRODUCTION

Hybridization and gene flow between taxa play a major role in evolution, acting as a force against divergence, and as a potential source of adaptive novelty (Abbott et al. 2013). Although identifying gene flow between species has been a long-standing problem in population genetics, the issue has received considerable recent attention with the analysis of shared ancestry between humans and Neanderthals (for example, Yang et al. 2012; Wall et al. 2013). With genomic data sets becoming available in a wide variety of other taxonomic groups, there is a need for reliable, computationally tractable methods that identify, quantify and date gene flow between species in large data sets.

A sensitive and widely used approach to test for gene flow is to fit coalescent models using maximum-likelihood or Bayesian methods (Pinho and Hey 2010). However, simulation and model fitting are computationally intensive tasks, and are not easily applied on a genomic scale. A simpler and more computationally efficient approach that is gaining in popularity is to test for an excess of shared derived variants using a four-taxon test (Kulathinal et al. 2009; Green et al. 2010; Durand et al. 2011). The test considers ancestral (‘A’) and derived (‘B’) alleles, and is based on the prediction that two particular SNP patterns, termed ‘ABBA’ and ‘BABA’ (see Methods), should be equally frequent under a scenario of incomplete lineage sorting without gene flow. An excess of ABBA patterns is indicative of gene flow between two of the taxa, and can be detected using Patterson’s *D* statistic (Green et al. 2010; Durand et al. 2011; see Methods for details). However, an excess of shared derived variants can arise from factors other than recent introgression, in particular non-random mating in the ancestral population due to population structure (Eriksson and Manica 2012). It is therefore important to make use of additional means to distinguish between these alternative hypotheses, for example, by examining the size of introgressed tracts (Wall et al. 2013), or the level of absolute divergence in introgressed regions (Smith and Kronforst 2013).

The *D* statistic was originally designed to be applied on a genome-wide or chromosome-wide scale, with block-jackknifing used to overcome the problem of non-independence between loci (Green et al. 2010). However, many researchers are interested in identifying particular genomic regions subject to gene flow, rather than simply estimating a genome-wide parameter. Theory predicts that the rate of gene flow should vary across the genome, both in the case of secondary contact after isolation (Barton and Gale 1993) as well as continuous gene flow during speciation (Wu 2001). Indeed, a maximum likelihood test for speciation with gene flow devised by Yang (2010) is based on detecting this underlying heterogeneity. Moreover, adaptive introgression might lead to highly localized signals of introgression, limited to the particular loci under selection.

Many methods for characterizing heterogeneity in patterns of introgression across the genome have been proposed. Several genomic studies have used *F_ST_* to characterize heterogeneity in divergence across the genome, often interpreting the variation in *F_ST_* as indicative of variation in rates of gene flow (for example, Hohenlohe et al. 2010). However, it is well established that, as a relative measure of divergence, *F_ST_* is dependent on within-population genetic diversity (Charlesworth 1998), and is therefore an unreliable indicator of how migration rates vary across the genome. In particular, heterogeneity in purifying selection and recombination rate could confound *F_ST_*-based studies (Noor et al. 2009, Hahn et al. 2012, Roesti et al. 2012, Cruickshank and Hahn, 2014). Various studies of admixture among human populations, or between humans and Neanderthals, have used probabilistic methods to assign ancestry to haplotypes, and infer how this ancestry changes across a chromosome (Sankararaman et al. 2008, Price et al. 2009, Henn et al. 2012, Lawson et al. 2012, Omberg et al. 2012, Maples et al. 2013, Churchhouse and Marchini 2013, Sankararaman et al. 2014). Other methods have modeled speciation with the allowance for variable introgression rates among loci (Garrigan et al. 2012, Roux et al. 2013), allowing the detection of more ancient gene flow.

There have also been recent attempts to characterize heterogeneity in patterns of introgression across the genome using the *D* statistic, calculated either in small windows (Smith and Kronforst 2013, Kronforst et al. 2013) or for individual SNPs (Rheindt et al. 2014). The robustness of the *D* statistic for detecting a genome-wide excess of shared derived alleles has been thoroughly explored (Green et al. 2010; Durand et al. 2011; Yang et al. 2012; Eaton and Ree 2013; Martin et al. 2013). However, it has not been established whether *D* provides a robust and unbiased means to identify individual loci with an excess of shared derived alleles, or to demonstrate that these loci have been subject to introgression. Any inherent biases of the *D* statistic when applied to specific loci have implications for methods that assume its robustness.

For example, Smith and Kronforst (2013) made use of the *D* statistic in a proposed test to distinguish between the hypotheses of introgression and shared ancestral variation at wing-patterning loci of *Heliconius* butterflies. Two wing-patterning loci are known to show an excess of shared derived alleles between co-mimetic populations of *Heliconius melpomene* and *Heliconius timareta* (*Heliconius* Genome Consortium, 2012). At one of these loci, phylogenetic evidence and patterns of linkage disequilibrium are consistent with recent gene flow (Pardo-Diaz et al. 2012). Nevertheless, Smith and Kronforst (2013) argue that this shared variation might represent an ancestral polymorphism that was maintained through the speciation event by balancing selection. Conceptually, this is not unlike the population structure argument of Eriksson and Manica (2012), except that here structure is limited to one or a few individual loci.

Smith and Kronforst proposed that the alternative explanations of introgression or ancestral polymorphism could be distinguished by considering absolute divergence within and outside of the loci of interest. Both hypotheses predict an excess of shared derived alleles at affected loci, but introgression should lead to reduced absolute divergence due to more recent coalescence at these loci, whereas the locus-specific population structure hypothesis predicts no reduction in absolute divergence at these loci compared to other loci in the genome. Loci with an excess of shared derived alleles, and therefore showing evidence of shared ancestry, were located by calculating the *D* statistic in non-overlapping 5 kb windows across genomic regions of interest, and identifying outliers using an arbitrary cutoff (the 10% of windows with the highest *D* values). The mean absolute genetic divergence (*d_XY_*) was then compared between the outliers and non-outliers, and found to be significantly lower in outlier windows, consistent with recent introgression (Smith and Kronforst 2013). This method makes two assumptions. First, that the *D* statistic can accurately identify regions that carry a significant excess of shared variation, and second, that *D* outliers do not have inherent biases leading to their co-occurrence with regions of low absolute divergence. These assumptions, which extend the use of *D* beyond its original definition, may be made by other researchers for similar purposes, but they remain to be tested.

Here, we first assess the reliability of the *D* statistic as a means to quantify introgression at individual loci. Using simulations of small sequence windows, we compare *D* to a related statistic that was developed by Green et al. (2010) specifically for estimating *f*, the proportion of the genome that has been shared, and we propose improvements to this statistic. We then use whole-genome data from several *Heliconius* species to investigate how these statistics perform on empirical data, and specifically how they are influenced by underlying heterogeneity in diversity across the genome. Lastly, we use a large range of simulated data sets to test the proposal that recent gene flow can be distinguished from shared ancestral variation based on absolute divergence in *D* outlier regions.

## RESULTS

### The *D* statistic is not an unbiased estimator of gene flow

Patterson’s *D* statistic was developed to detect, but not to quantify introgression. We used the deterministic derivation of the expected value of *D* (*E*[*D*]) provided by Durand et al. (2011, Equation 5) to test how sensitive the value of *D* is to other factors apart from the proportion of introgression. We define the proportion of introgression (*f*) as the proportion of haplotypes in the recipient population (P_2_) that trace their ancestry through the donor population (P_3_) at the time of gene flow (Figure 1A). The expected *D* value increases with the proportion of introgression (*f*), but not linearly (Figure 1B) and expected *D* increases as population size decreases (Figure 1B,C). The split times between populations also have a small effect, with a more recent split between P_1_ and P_2_ leading to higher expected values of *D* (Figure 1C). This implies that empirically calculated values of the *D* statistic will depend on various parameters other than the amount of gene flow, irrespective of the number of sites analyzed.

**Figure 1.**
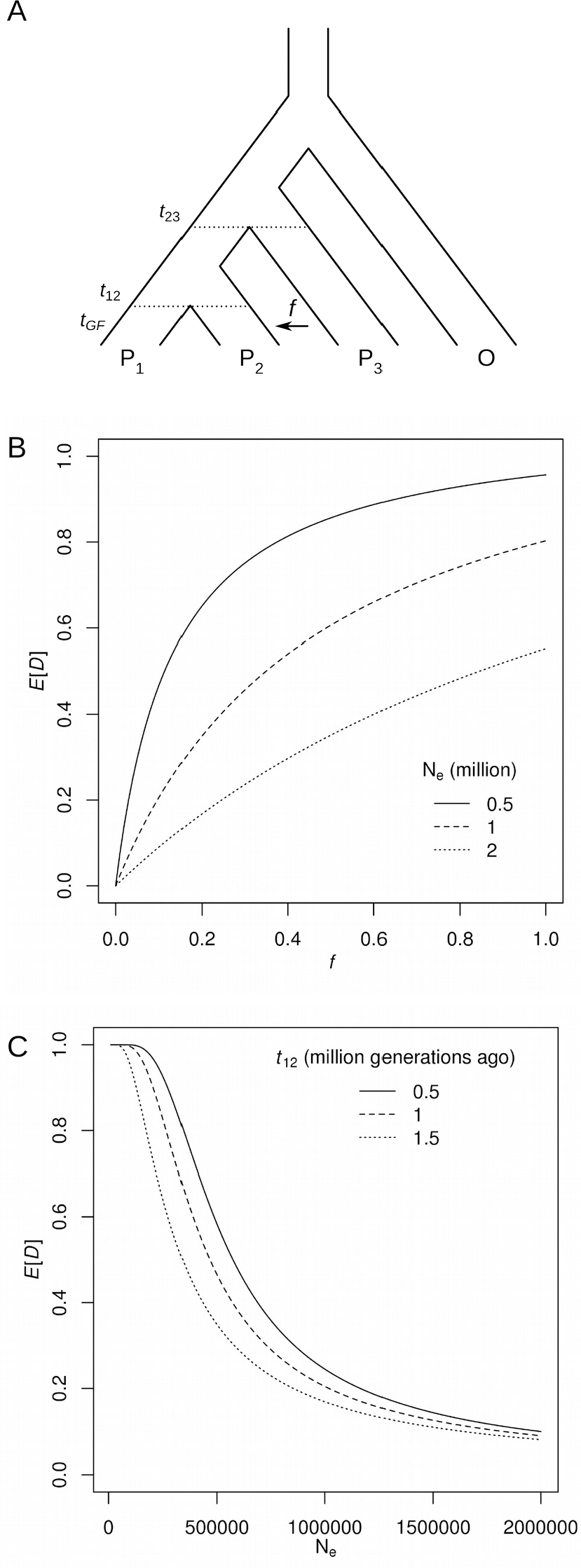
Expected value of the *D* statistic. **A.** Durand et al. (2011)’s derivation of the expected value of Patterson’*s D* statistic *E*[*D*] depends on the two split times, *t_12_* and *t_23_*, separating populations P_1_, P_2_ and P_3_. It assumes a single instantaneous admixture event from P_3_ to P_2_ at *t*_*G*F_, after which a proportion *f* of P_2_ individuals trace their ancestry through P_3_. The effective population size, *N_e_*, is constant through time and the same in all populations. **B.** The expected value of *D* as a function of *f*, the proportion of introgression, at three different effective population sizes: 0.5, 1 and 2 million. Split times are fixed at 1 million generations for *t_12_* and 2 million generations for *t_23_*. **C.** The expected value of *D* as a function of *N_e_*, showing the effect of varying t_12_. In all three cases, *t_23_* is set at 2 million generations ago, and *f* is set to 0.1.

### Direct estimators of *f* outperform *D* on simulated data

Analysis of simulated data confirmed that the *D* statistic is not an appropriate measure for quantifying introgression over small genomic windows, but that direct estimation of the proportion of introgression (*f*) provides a more robust alternative. The *D* statistic (Equation 1, Methods) was compared to the *f* estimator of Green et al. (2010) (Equation 4, Methods), which is referred to as 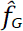 below, along with two proposed modified versions of this statistic (Equations 5 and 6, Methods). The first, 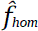 (Equation 5), is similar to 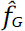 in that it explicitly assumes unidirectional gene flow from P_3_ to P_2_, but makes a further assumption that maximal introgression would lead to complete homogenization of allele frequencies in P_2_ and P_3_. This is a conservative assumption, as an extremely high rate of migration would be necessary to attain a maximal value of 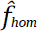. The second, 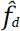 (Equation 6), is dynamic in that it allows for bidirectional introgression on a site-by-site basis, setting the donor population at each site as that which has the higher frequency of the derived allele. These *f* estimators are distinct from the *F_2_*, *F_3_* and *F_4_* statistics of Reich et al. (2009, 2012) and Patterson et al. (2012), which all test for correlated allele frequencies associated with introgression (much like the *D* statistic). However, Patterson’s (2012) *F_4_*-ratio is conceptually very similar to the *f* estimators discussed here, in that it estimates the proportional contribution of a donor population.

To compare the utility of these statistics for quantifying introgression in small genomic windows, we simulated sequences from four populations: P_1_, P_2_ and P_3_ and outgroup O, with the relationship (((P_1_,P_2_),P_3_),O), with a single instantaneous gene flow event, either from P_3_ to P_2_ or from P_2_ to P_3_. Simulations were performed over a range of different values of *f* (the probability that any particular haplotype is shared during the introgression event), and with various window sizes, recombination rates and times of gene flow.

A subset of the results are shown in Figure 2, and full results are provided in Figure S1. In general, the *D* statistic proved sensitive to the occurrence of introgression, with strongly positive values for any non-zero value of *f*. However, it was a poor estimator of the amount of introgression, as defined by the simulated value of *f* (Figure 2). Moreover, *D* values showed dramatic variance, particularly at low simulated values of *f*. Even in the absence of any gene flow, a considerable proportion of windows had intermediate *D* values. This variance decreased with increasing window size and recombination rate (Figure S1).

**Figure 2.**
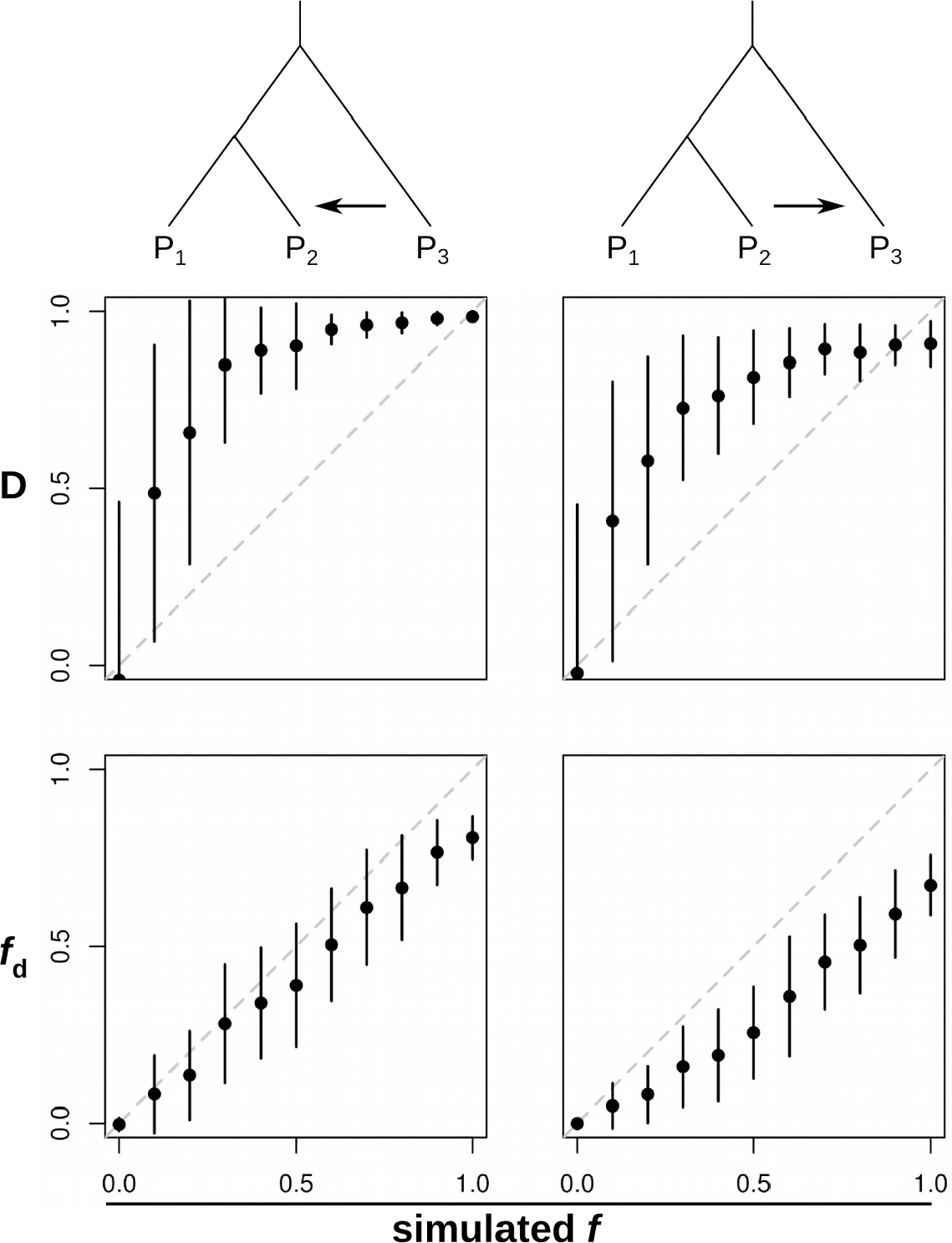
Comparing statistics to detect and quantify introgression. Results from a subset of the simulations: window size 5 kb, time of gene flow (*t_GF_*) 0.1 × 4*N* generations ago, and population recombination rate 0.01. See Figures S1A-I for full results. Plots show means and standard deviations for *D* and *f̂_d_*, calculated over 100 simulated sequences (see Methods for details). Simulations covered 11 different values of *f,* the proportion of introgression. Gene flow was simulated either from P_3_ to P_2_ (left-hand column) or from P_2_ to P_3_ (right-hand column). Dashed diagonal lines show the expectation of a perfect estimator of *f*.

In simulations of gene flow from P_3_ to P_2_, all three *f* estimators gave fairly accurate estimates of the simulated *f* value, provided gene flow was recent (Figures 2, S1A). When gene flow occurred further back in time, *f* estimators tended to give underestimates, but were nevertheless well correlated with the simulated *f* value (Figure S1A). This is unsurprising, as genetic drift and the accumulation of mutations in these lineages after the introgression event should dilute the signal of introgression. In simulations of gene flow in the opposite direction, from P_2_ to P_3_, both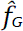 and 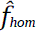 showed considerable stochasticity, particularly when recombination rates were low and gene flow was recent (Figure S1). The size of the window had little effect on this behavior (Figure S1), implying that it was not an effect of the number of sites analyzed, but rather the level of independence among sites. This is also to be expected, as these statistics can give values greater than 1 where derived allele frequencies happen by chance to be higher in P_3_ than P_2_ (see Methods). Unlike these two statistics, 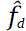 behaved predictably at all recombination rates and times of gene flow, giving estimates that were fairly well correlated with the simulated *f*, but underestimating its absolute value (Figure 2, Figure S1).

Generally, the variance in 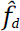 was lower than in the other two *f* estimators (Figure S1A-I). Unlike the *D* statistic, 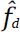 displayed minimal variance at low simulated values of *f* (Figure 2). However, all four statistics showed greater variance and more extreme values when recombination rates were lower (Figure S2), as expected given that decreased recombination reduces the number of independent sites analyzed.

Although none of the examined measures were able to accurately quantify both forms of introgression in all cases, 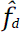 has some appealing characteristics as a measure to identify introgressed loci in a genome scan approach. It has low variance and is not prone to false positives when gene flow is absent and recombination rare. In all of our simulations, it provided estimates that were proportional to the simulated level of introgression. Although it tended toward underestimates, genome scans for introgressed loci would primarily be interested in relative rates of introgression across the genome, rather than absolute rates.

### *f* estimators are robust to variation in nucleotide diversity across the genome

Analysis of published whole-genome data from *Heliconius* species confirmed that Patterson’s *D* statistic was prone to extreme values in regions of low diversity, whereas *f* estimators were not (Figure 3A-C). We re-analyzed published whole-genome sequence data from two closely related *Heliconius* butterfly species, *Heliconius melpomene* and *Heliconius timareta*, and four outgroup species from the related silvaniform clade. The races *H. melpomene amaryllis* and *H. timareta thelxinoe* are sympatric in Peru, and show genome-wide evidence of gene flow (Martin et al. 2013), with particularly strong signals at two wing-patterning loci: *HmB*, which controls red pattern elements, and *HmYb,* which controls yellow and white pattern elements (*Heliconius* Genome Consortium 2012, Pardo-Diaz et al. 2012). To determine whether heterogeneity in diversity across the genome may influence *D* and the *f* estimators, we calculated *D*, 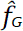,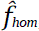, 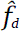 and nucleotide diversity (π) in non-overlapping 5 kb windows across the genome. Variance in the *D* statistic was highest among windows with low nucleotide diversity and decreased rapidly with increasing diversity (Figure 3A, Figure S4). Windows from the wing-patterning loci were among those with the highest *D* values, but there were many additional windows with *D* values approaching or equal to 1. By contrast, 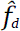, calculated for all windows with positive *D*, was far less sensitive to the level of diversity, with most outlying windows showing intermediate levels of diversity (Figure 3B). Notable exceptions were windows located within the wing-patterning regions, which tended to have high 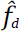 values and below average diversity. This is consistent with the strong selection known to act upon the patterning loci. The lack of extreme 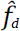 values in windows with low diversity suggests that most of the *D* outliers are spurious, and that 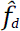 provides a better measure of whether a locus has shared ancestry between species. Finally, we also tested the other two *f* estimators described here: 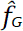 and 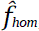 (Equations 4 and 5, Methods). Both performed similarly to 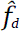 except that both had higher variance (Figures S3, S4), consistent with the simulations reported above (Figure S1), and both gave a considerable number of values greater than 1, confirming that 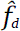 was the most conservative and stable statistic.

**Figure 3.**
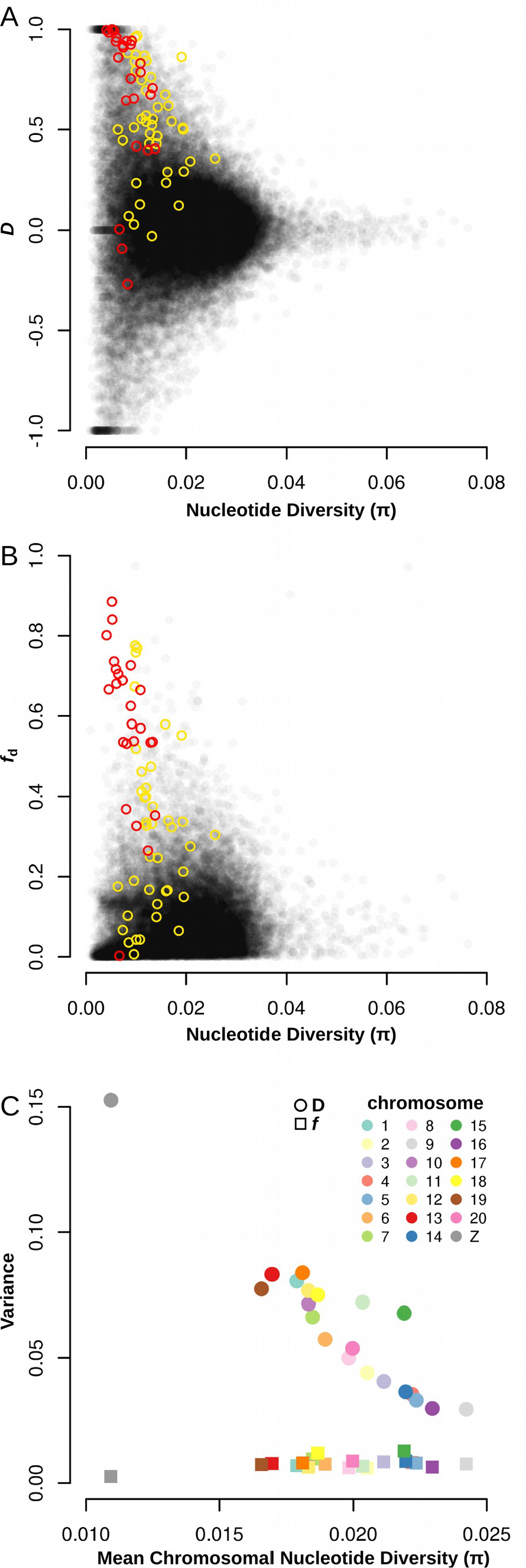
Effects of genetic diversity on the *D* and *f_d_* in *Heliconius* whole genome data. **A,B.** Values of *D* and 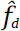 for non-overlapping 5 kb windows across the genome, plotted against nucleotide diversity. *f̂_d_* values are only plotted for windows with D ≥ 0. Data from Martin et al. 2013. Taxa used are as follows, P_1_: *Heliconius melpomene aglaope*, P_2_: *Heliconius melpomene amaryllis*, P_3_: *Heliconius timareta thelxinoe*, O: four *Heliconius* species from the silvaniform clade (see Methods for list). Colored points show windows located within the wing-patterning loci *HmB* (red) and *HmYb* (yellow). **C.** The variance among *D* and *f_d_* values for each chromosome, plotted against the mean nucleotide diversity from all windows for each chromosome.

Taken together, these findings demonstrate that, when small genomic windows are analyzed, a high *D* value alone is not sufficient evidence for introgression. Many of the *D* outlier loci probably represent statistical noise, concentrated in regions of low diversity, whereas outliers for the *f* estimates, and particularly 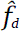, tend to be less biased.

This effect could also be observed on the scale of whole chromosomes. The variance in *D* among 5 kb windows for each of the *Heliconius* chromosomes (n = 21) was strongly negatively correlated with the average diversity per chromosome (r(19)=-0.936, p < 0.001; Figure 3C). This relationship was most clearly illustrated by the Z chromosome: it had the lowest diversity by some margin, as expected given its reduced effective population size, and the highest variance among *D* values for 5 kb windows, despite the fact that previous chromosome-wide analyses suggest very limited gene flow affecting this chromosome (Martin et al. 2013). By contrast, the variance in 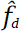, estimated for all windows with positive *D*, had a weak positive correlation with the mean diversity per chromosome (r(19) = 0.440, p < 0.05). This was driven by the fact that the Z chromosome had the lowest diversity and also the lowest variance in 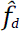, as expected given the reduced gene flow affecting this chromosome. When the Z chromosome was excluded, there was no significant relationship between the variance in 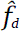 values and average diversity (r(18) = 0.092, p > 0.05). We also considered the effect of window size on the variance of *D* and the estimators of *f*. As window size increases, the higher variance of *D* in regions of lower diversity persists, but becomes less extreme (Figure S4). In summary, these data show that extreme *D* values, both positive and negative, occur disproportionately in genomic regions with lower diversity, whereas 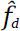 values are less biased by underlying heterogeneity in genetic variation.

### Inherent biases in the *D* and *f̂* statistics confound a test to distinguish between introgression and shared ancestral variation

The biases associated with the *D* statistic described above may have important consequences for methods that use *D* to identify candidate introgressed regions. For example, Smith and Kronforst (2013) proposed a method to discriminate between gene flow and shared ancestral variation that relies upon *D* values calculated for small genomic regions (see Introduction). Briefly, the Smith and Kronforst test calculated *D* for all non-overlapping 5 kb windows. Absolute divergence (*d_XY_*) was then compared between the set of windows that were outliers for the *D* statistic (defined as the windows with the top 10% of *D* values) and the remaining 90% of non-outlier windows. The method predicts that introgression between species at a specific genomic region should reduce the between-species divergence in this region as compared to the rest of the genome, while shared ancestry due to ancestral population structure would not lead to lower divergence. We first confirmed this prediction using simulations, and then assessed whether biases in the *D* statistic might affect the power of the method.

To test the prediction that introgression and ancestral population structure leave distinct footprints in terms of absolute divergence, 10 000 sequence windows for three populations (P_1_,P_2_,P_3_) and an outgroup (O) were simulated. 9000 windows were defined as ‘Background’, having the topology (((P_1_,P_2_),P_3_),O), without any gene flow or population structure. The remaining 1000 windows were defined as ‘Alternate’ and were subject to either gene flow or structure (see Methods for details). Ten percent of windows were defined as Alternate to match Smith and Kronforst’s design, wherein the top 10% of *D* values are taken as outliers. Three different Alternate scenarios were considered: gene flow from P_2_ to P_3_, gene flow from P_3_ to P_2_ and ancestral structure leading to shared ancestry between P_2_ and P_3_. For all scenarios, Alternate windows were defined with the topology ((P_1_,(P_2_,P_3_)),O). In the gene flow scenarios, the split time between P_2_ and P_3_ in the Alternate topology was set to be more recent than the split between P_1_ and P_2_ in the Background topology (Figure 4A,B). In the ancestral structure scenario, the split time between P_1_ and P_2_ in the Alternate topology was set to be more ancient than the split between P_2_ and P_3_ in the Background topology (Figure 4C). This was designed to model a region of the genome undergoing balancing selection or some other process that maintains polymorphism at particular loci prior to the speciation event. Gene flow or structure in the Alternate windows can be considered to be complete (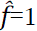). For example, under gene flow from P_2_ to P_3_, all P_3_ alleles trace their ancestry through P_2_ at the time of gene flow. This simplified design, where gene flow or structure is absent in 90% of the sequences and complete in 10%, allowed for the most straightforward and predictable test of Smith and Kronforst’s method; if the logic of the method does not follow in this extreme scenario, it is unlikely to do so in more complex situations.

**Figure 4.**
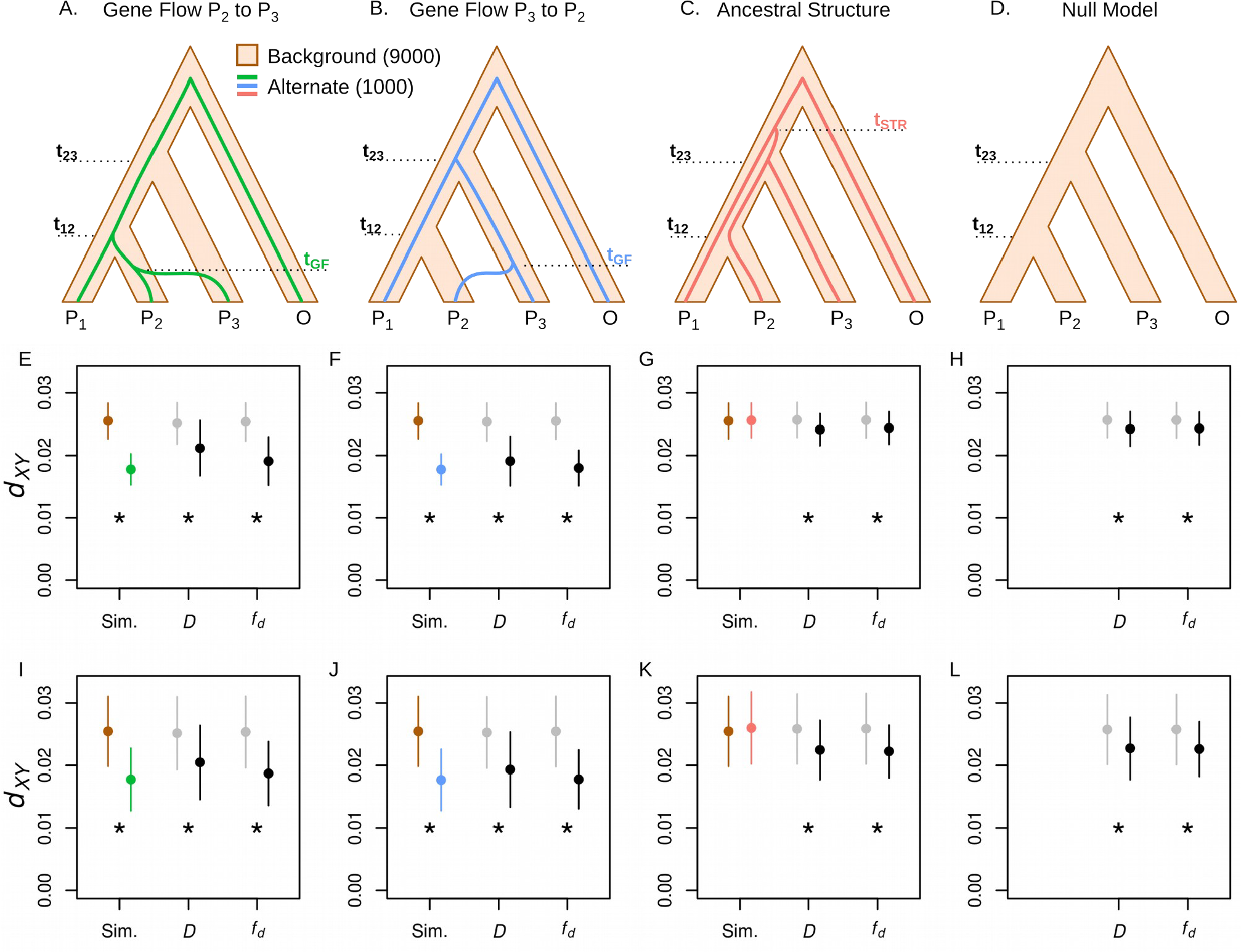
Simulations to evaluate a method to distinguish introgression from shared ancestral variation. **A-C.** Combined models were made up of 9000 sequence windows simulated under the “Background” topology (brown outline) and 1000 windows simulated under an “Alternate” topology (colored line). Three distinct evolutionary scenarios were simulated by varying the split times *t_12_*, *t_23_*, *t_GF_* and *t_STR_*; **A, E, I**: gene flow from P_2_ to P_3_, **B, F, J**: gene flow from P_3_ to P_2_, **C, G, K**: ancestral structure. **D, H, L.** Null models were made up of 10 000 sequences simulated under the Background topology only. **E-L.** Example data from a single simulated dataset for each of the four types of models. Split times (in units of 4*N* generations) were as follows: *t_12_* = 0.6 in all four cases, *t_23_* = 0.8 in all four cases, *t_GF_* = 0.4 in both gene flow models and *t_STR_* = 1.0. Points show mean and standard deviation for P_2_-P_3_ *d_XY_* calculated over subsets of trees: simulated Background and Alternate trees (brown and colored points) or non-outliers and outliers (gray and black points) identified using the *D* and 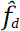 statistics. A significant reduction in P_2_-P_3_ *d_XY_* for the Alternate compared to Background windows, or for outliers compared to non-outliers, is indicated by astrices. **E-H** show results of simulations with a population recombination rate (4Nr) of 0.01. **I-L** show results for the same models, but with a population recombination rate (4Nr) of 0.001.

For each of the three evolutionary scenarios, 120 different permutations of split times and times of gene flow or structure were simulated. The split times and times of gene flow for all models are given in the first three columns of Tables S1, S2 and S3. To simplify our comparisons between models, we focused specifically on *d_XY_* between P_2_ and P_3_, the most relevant parameter when testing for introgression between P_2_ and P_3_. We tested whether P_2_-P_3_ *d_XY_* was significantly lower in the Alternate windows (those that had experienced gene flow or structure) compared to the Background windows, using a Wilcoxon rank-sum test, with Bonferroni correction over all 120 models of the same type, and a significance threshold of 99%. We first performed simulations with a recombination rate parameter (4Nr) of 0.01, and later repeated all simulations at 4Nr = 0.001. The results of all tests are given in full in Tables S1, S2 and S3. These results are summarized in Table 1 and Figure 5, and a single illustrative example for each model type is given in Figure 4E-L.

**Figure 5.**
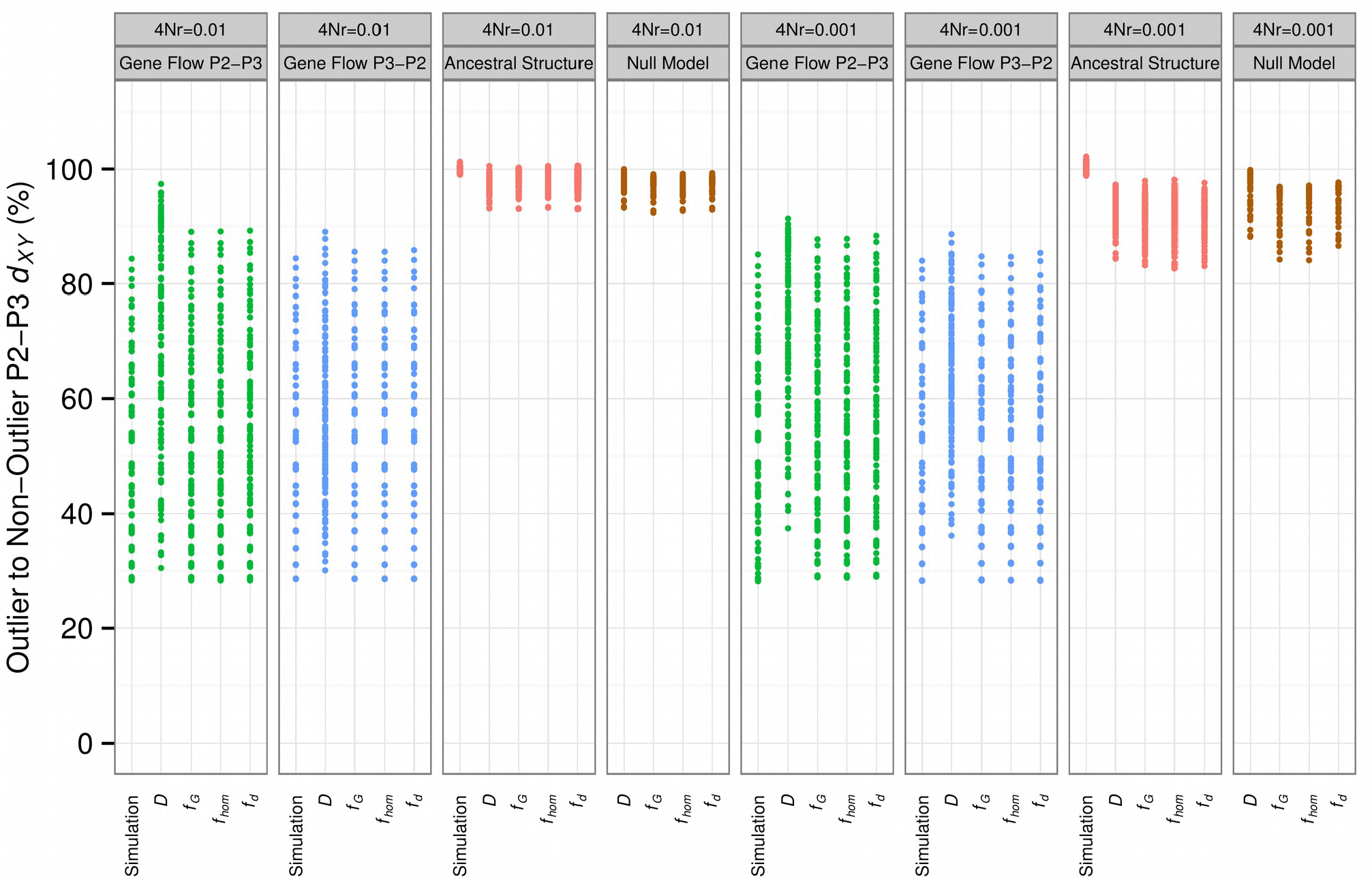
Mean *d_XY_* between P_2_ and P_3_ in outlier windows as a percentage of P_2_-P_3_ *d_XY_* in non-outlier windows. Outlier windows defined by Alternate or Background topology (Simulation) or by outlying *D* and *f̂* values, as per Figure 4. Model types shown in color (gene flow from P_2_ to P_3_, green; gene flow from P_3_ to P_2_, blue; ancestral structure, red; null model, brown). Results for two different recombination rates are shown (4Nr = 0.01, left; 4Nr = 0.001, right).

**Table 1.** Number of simulated models in which P_2_-P_3_ *d_XY_* is significantly reduced in Alternate vs Background windows, or in outlier vs non-outlier windows for the introgression statistics

| Model Type<br>(No. of models) | Number of models with significantly reduced mean $P_2$-$P_3$ $d_{XY}$ | | | | |
| --- | --- | --- | --- | --- | --- |
| | Alternate vs<br>Background | $D$ outliers vs<br>non-outliers | $f_d$ outliers vs<br>non-outliers | $f_G$ outliers vs<br>non-outliers | $f_{hom}$ outliers vs<br>non-outliers |
| 4Nr = 0.01 |  |  |  |  |  |
| Gene Flow $P_3 \rightarrow P_2$<br>(120) | 120 | 119 | 120 | 120 | 120 |
| Gene Flow $P_2 \rightarrow P_3$<br>(120) | 120 | 120 | 120 | 120 | 120 |
| Ancestral Structure<br>(120) | 0 | 105 | 72 | 67 | 66 |
| Null<br>(45) | 0 | 39 | 44 | 45 | 44 |
| 4Nr = 0.001 |  |  |  |  |  |
| Gene Flow $P_3 \rightarrow P_2$<br>(120) | 120 | 120 | 120 | 120 | 120 |
| Gene Flow $P_2 \rightarrow P_3$<br>(120) | 120 | 120 | 120 | 120 | 120 |
| Ancestral Structure<br>(120) | 0 | 120 | 119 | 118 | 118 |
| Null<br>(45) | 0 | 32 | 45 | 45 | 45 |

Significantly lower $d_{XY}$ was evaluated using a Wilcoxon rank sum test, with a 99% significance threshold after Bonferroni correction over the 120 models of each type (except for Null models, of which there were 45).

As predicted, in all models simulating gene flow, average *d_XY_* between P_2_ and P_3_ was significantly lower in Alternate windows compared to Background windows. In contrast, in all models simulating ancestral population structure, there was no significant difference in P_2_-P_3_ *d_XY_* between the background and alternate windows, again in agreement with predictions. These findings therefore demonstrate that the intuitive premise of Smith and Kronforst’s (2013) method is justified.

We then tested whether introgression could be distinguished from shared ancestral variation where loci with shared ancestry are not known (as would be the situation with empirical data), but are instead inferred by selecting the top 10% of *D* values (outliers), following the Smith and Kronforst method. We also tested this method using the top 10% of *f* estimates among windows with positive *D* (using 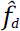, 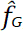 and 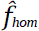). Using the *D* statistic to identify outliers, mean *d_XY_* between P_2_ and P_3_ was significantly reduced in outlier windows as compared to non-outlier windows in all 120 models simulating gene flow from P_3_ to P_2_, and all but one of the models simulating gene flow from P_2_ to P_3_. The single non-significant case had the most ancient possible *t_23_* and the most recent possible *t_12_*, and only 11.9% of *D* outlier windows were genuine Alternate windows, the lowest recall of any model. Using any of the three *f* estimators, mean *d_XY_* between P_2_ and P_3_ was significantly reduced in outlier windows in all gene flow models.

However, mean P_2_-P_3_ *d_XY_* was also significantly reduced in *D* and ***f*** outlier windows in more than half of the 120 models simulating ancestral population structure (Figure 4C, 4G, 5; Tables 1, S3). This demonstrates that a simple test for reduced divergence in P_2_-P_3_ *d*_XY_ among *D* or 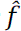 outlier windows would, under a range of ancestral structure scenarios, produce results consistent with introgression. The fact that this bias was similar whether *D* or *f* estimators were used to identify outliers indicates that there is an inherent tendency in all of these statistics toward regions with below-average divergence between P_2_ and P_3_. To confirm this finding, we analyzed a set of simulations using a null model, with no gene flow or structure in any of the 10 000 windows, over 45 permutations of split times (Table S4). Outlier windows showed significantly reduced *d_XY_* between P_2_ and P_3_ in most or all of the null models. Finally, we repeated all of these simulations with a lower within-window recombination rate parameter (4Nr) of 0.001. This tended to increase the reduction in P_2_-P_3_ *d_XY_* for outliers in ancestral structure and null models (Figures 4I-L, 5), with at most 3 of the ancestral structure models showing non-significant drops in P_2_-P_3_ *d_XY_* for outliers, and most or all null models showing significantly lower P_2_-P_3_ *d_XY_* for outliers, regardless of the statistic used (Table 1).

In summary, although shared ancestral variation and introgression can theoretically be distinguished based on the fact that only the latter should reduce *d_XY_* between P_2_ and P_3_, an inherent bias in both the *D* and ***f*** statistics makes a simple test for a statistical difference in *d_XY_* between outliers and non-outliers problematic. Both *D* and ***f*** outliers tended toward windows with lower P_2_-P_3_ *d_XY_*, regardless of the underlying evolutionary history, and particularly when recombination rates were low. In the absence of any gene flow, the outliers must therefore be identifying windows that coalesce more recently in the ancestral population. However, even when the reduction in P_2_-P_3_ *d_XY_* was significant for ancestral structure or null models, it was typically smaller than the reductions in *d_XY_* seen in the gene flow models (Figure 5). In the presence of gene flow, some windows coalesce more recently than the species split, so the magnitude of the reduction in P_2_-P_3_ *d_XY_* is greater. This difference could potentially be used to distinguish introgression from shared ancestral variation, but can not be done with a simple significance test, and will require a more sophisticated model-fitting approach.

## DISCUSSION

With the advent of population genomics, studies of species divergence have moved from simply documenting inter-specific gene flow, towards the identification of specific genomic regions that show strong signals of either introgression or divergence (The *Heliconius* Genome Consortium, 2012; Garrigan et al. 2012; Staubach et al. 2012; Roux et al. 2013, Bosse et al. 2014, Huerta-Sánchez et al. 2014, Sankararaman et al. 2014). This is a useful goal for many reasons. It can permit the identification of large-scale trends, such as chromosomal differences, and the fine-scale localization of putative targets of adaptive introgression for further characterization. Therefore, simple and easily computable statistics that can be used to identify loci with a history of introgression have considerable appeal.

Previous studies have explored the behavior of Patterson’s *D* statistic, a test for gene flow based on detecting an inequality in the numbers of ABBA and BABA patterns, using whole genome analyses across large numbers of informative sites (Green et al. 2010; Yang et al. 2012; Eaton and Ree 2013; Martin et al. 2013; Wall et al. 2013). These studies have shown that *D* is a robust method when applied as intended: to test for an excess of shared variation on a genome-wide scale. Indeed a major strength of the ABBA-BABA test is that it combines data from across the genome, accounting for chance fluctuations among loci, and therefore is able to detect the net effect of gene flow. Moreover, the non-independence among linked sites can be accounted for by block-jackknifing (Green et al. 2010).

However, it is not clear whether *D* can be extended beyond its original use to identify specific loci with introgressed variation. We have documented two main problems with this approach. Firstly, *D* is not an unbiased estimator of the amount of introgression that has occurred. In particular it is influenced by effective population size (*N_e_*), leading to more extreme values when *N_e_* is low. Secondly, when calculated over small windows, it is highly stochastic, particularly in genomic regions of low diversity and low recombination rate, such that *D* outliers will tend to be clustered within these regions. Local reductions in genetic diversity along a chromosome can come about through neutral processes, such as population bottlenecks, but also through directional selection. Therefore, these problems may be exacerbated in studies specifically interested in loci that experience strong selective pressures, as this would increase the likelihood of detecting chance outliers at such loci.

Direct estimation of *f,* the proportion of introgression, holds more promise as a robust method for detecting introgressed loci. Green et al. (2010) proposed that *f* could be estimated by comparing the observed difference in the number of ABBA and BABA patterns to that which would be expected in the event of complete introgression. As this expected value is calculated from the observed data, this method controls for differences in the level of standing variation, making it more suitable for application to small regions. In Green et al.’s approach, complete introgression from P_3_ to P_2_ was taken to mean that P_2_ would come to resemble a subpopulation of lineage P_3_. Here we make the conservative assumption that complete introgression would lead to homogenization of allele frequencies, such that the frequency of the derived allele in P_2_ would be identical to that in P_3_. Green et al.’s approach assumed unidirectional introgression from P_3_ to P_2_, but can lead to spurious values when introgression occurs in the opposite direction. We have therefore proposed a new dynamic estimator of *f,* in which the donor population can differ between sites, and is always the population with the higher frequency of the derived allele. Although this conservative estimator leads to slight underestimation of the amount of introgression that has occurred, it provides an estimate that is roughly proportional to the level of introgression, regardless of the direction. It is therefore a more suitable measure for identifying introgressed loci. This is supported by our analysis of whole-genome data from *Heliconius* butterflies, where many 5 kb windows had maximal *D* values (*D* = 1), but only a few had high 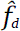 values, the vast majority of which were located around the wing-patterning loci previously identified as being shared between these species through adaptive introgression (*Heliconius* Genome Consortium 2012, Pardo-Diaz et al. 2012).

The sensitivity of *D* to heterogeneous genomic diversity is likely to affect studies that have drawn conclusions from *D* statistics calculated for particular genome regions. For example, Wall et al. (2013) identified regions carrying putative long (8-100kb) haplotypes segregating in European humans, and then found that these regions showed evidence of a Neanderthal origin, as indicated by elevated *D* statistics. However, it may be that such haplotypes would be over-represented in low-recombination regions, which also tend to have reduced diversity in humans and many other species (Cutter and Payseur 2013). In another recent *Heliconius study*, *F_ST_* was calculated for 5kb windows across the genome. Windows showing increased differentiation between *H. melpomene* and *H. pachinus* (according to *F_ST_*) also showed significantly elevated *D* statistics in a test for introgression between the same species pair (Kronforst et al. 2013). This illustrates how the sensitivity of both *D* and *F_ST_* to within-species diversity can produce conflicting results. This sensitivity is likely to be particularly problematic in studies using very small genomic regions. At the extreme, Rheindt et al. (2014) calculated *D* for single SNPs and predicted that genes linked to SNPs with outlying *D* values are more likely to have been introgressed.

In the present study, to investigate whether biases in *D* when calculated over small regions could influence subsequent analyses, we investigated a recently proposed method to distinguish between introgression and shared ancestral variation (Smith and Kronforst 2013). The premise of this test is that introgression should result in an excess of shared derived alleles and a reduction in absolute divergence (*d_XY_*), whereas shared ancestral variation will exhibit the former but not the latter signature. Our simulations confirmed that the intuitive predictions of this method are valid, but also showed that this test can be misled by the use of *D* to identify outliers. Windows that were outliers for *D* exhibited below average *d_XY_* in simulations with gene flow, but also in most simulations with ancestral structure, or where both gene flow and ancestral population structure were absent. All three *f* estimators also failed to distinguish between introgression and ancestral structure in many models. This implies that all of these statistics are systematically biased toward regions that coalesce more recently, regardless of whether gene flow has occurred.

We predict that *D* would have additional problems in real genomes, where selective constraint leads to a correlation between within-species diversity and between-species divergence, causing *D* outliers to be even more strongly associated with reduced *d_XY_*. However, it is notable that the reduction in divergence among *D* and ***f*** outliers was almost always greater in simulations with introgression than in simulations with ancestral structure or with no alternate topology, across a large range of split times and dates of gene flow. There may, therefore, be considerable information about the evolutionary history of DNA sequences present in the joint distribution of *d_XY_* and 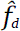. On the other hand, in real data, levels of divergence can vary dramatically due to heterogeneity in selective constraint, mutation rate and recombination rate, which would exaggerate the problems described here. Even an unbiased statistic, when applied to small genomic windows, would be confounded by heterogeneity in recombination rate across the genome. In regions of reduced recombination, fewer independent data points are sampled by each window, so extreme estimates become more likely. Heterogeneity in recombination rate is therefore an essential consideration in any study that aims to scan the genome for regions of interest.

### Conclusions

In an era of increasing availability of genomic data, there is a demand for simple summary statistics that can reliably identify genomic regions that have been subject to selection, introgression and other evolutionary processes. It seems unlikely, however, that any single summary statistic will be able to reliably distinguish these processes from noise introduced by demography, drift and heterogeneity in recombination rate. Here we have shown that, while Patterson’s *D* statistic provides a robust signal of shared ancestry across the genome, it should not be used for naïve scans to ascribe shared ancestry to small genomic regions, due to its tendency toward extreme values in regions of reduced variation. Estimation of *f*, the proportion of introgression, particularly using our proposed statistic 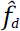, provides a better means of identifying putatively introgressed regions. Nevertheless, both *D* and 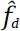 tend to identify regions of reduced inter-species divergence, even in the absence of gene flow, which may confound tests to distinguish between recent introgression and shared ancestral variation based on absolute divergence (*d_XY_*) in outlier regions. However, the joint distribution of *d_XY_* and 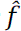 statistics may be a useful summary statistic for model-fitting approaches to distinguish between these evolutionary hypotheses.

## MATERIAL AND METHODS

### Statistics used to detect shared ancestry

In this study, we focused on an approach to identify an excess of shared derived polymorphisms, indicated by the relative abundance of two SNP patterns termed “ABBAs” and “BABAs” (Green et al. 2010). Given three populations and an outgroup with the relationship (((P_1_, P_2_), P_3_), O) (Figure 1A), ABBAs are sites at which the derived allele “B” is shared between the non-sister taxa P_2_ and P_3_, while P_1_ carries the ancestral allele, as defined by the outgroup. Similarly, BABAs are sites at which the derived allele is shared between P_1_ and P_3_, while P_2_ carries the ancestral allele. Under a neutral coalescent model, both patterns can only result from incomplete lineage sorting or recurrent mutation, and should be equally abundant in the genome (Durand et al. 2011). A significant excess of ABBAs over BABAs is indicative either of gene flow between P_2_ and P_3_, or some form of non-random mating or structure in the population ancestral to P_1_, P_2_ and P_3_. This excess can be tested for, using Patterson’s *D* statistic,

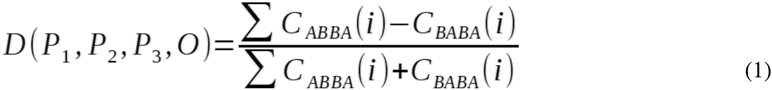

where *C_ABBA_(i)* and *C_BABA_(i)* are counts of either 1 or 0, depending on whether or not the specified pattern (ABBA or BABA) is observed at site *i* in the genome. Under the null hypothesis of no gene flow and random mating in the ancestral population, *D* will approach zero, regardless of differences in effective population sizes (Durand et al. 2011). Hence, a *D* significantly greater than zero is indicative of a significant excess of shared derived alleles between P_2_ and P_3_.

If population samples are used, then rather than binary counts of fixed ABBA and BABA sites, the frequency of the derived allele at each site in each population can be used (Green et al. 2010, Durand et al. 2011), effectively weighting each segregating site according to its fit to the ABBA or BABA pattern, with

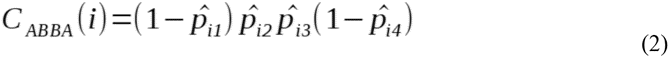

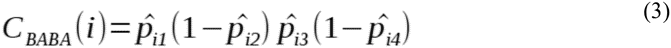

where *p_ij_* is the frequency of the derived allele at site *i* in population *j*. These values are then used in equation 1 to calculate *D* (Durand et al. 2011).

Green et al. (2010) also proposed a related method to estimate *f*, the fraction of the genome shared through introgression (Green et al. 2010, Durand et al. 2011). This method makes use of the numerator of equation 1, the difference between sums of ABBAs and BABAs, which is called *S*. In the example described above, with ((P_1_,P_2_),P_3_),O), the proportion of the genome that has been shared between P_2_ and P_3_ subsequent to the split between P_1_ and P_2_ can be estimated by comparing the observed value of *S* to a value estimated under a scenario of complete introgression from P_3_ to P_2_. P_2_ would then resemble a lineage of the P_3_ taxon, and so the denominator of equation 1 can be estimated by replacing P_2_ in equations 2 and 3 with a second lineage sampled from P_3_, or by splitting the P_3_ sample into two,

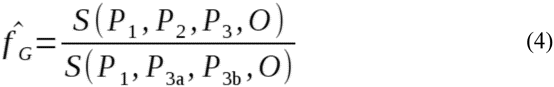

where P_3a_ and P_3b_ are the two lineages sampled from P_3_. Splitting P_3_ arbitrarily in this way may lead to stochastic errors at individual sites, particularly with small sample sizes. These should be negligible when whole-genome data are analyzed but could easily lead to erroneous values of 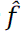(including 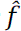 > 1) when small genomic windows are analyzed, as in the present study. We therefore used a more conservative version, in which we assume that complete introgression from P_3_ to P_2_ would lead to complete homogenization of allele frequencies. Hence, in the denominator, P_3a_ and P_3b_ are both substituted by P_3_:

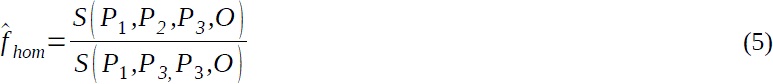

While this conservative assumption may lead to underestimation of the proportion of sites shared, it also reduces the rate of stochastic error. Moreover, in the present study, we are less concerned with the absolute value of 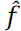, and more with the relative values of 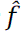 between genomic regions.

The 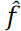 statistic assumes unidirectional gene flow from P_3_ to P_2_ (i.e. P_3_ is the donor and P_2_ is the recipient). Since the branch leading to P_3_ is longer than that leading to P_2_ (Figure 1A), gene flow in the opposite direction (P_2_ to P_3_) is likely to generate fewer ABBAs. Thus, in the presence of gene flow from P_2_ to P_3_, or in both directions, the 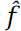 equation should lead to an underestimate. However, when small genomic windows are analyzed, the assumption of unidirectional gene flow could lead to overestimates, because any region in which derived alleles are present in both P_2_ and P_3_, but happen to be at higher frequency in P_2_, will yield *f* estimates that are greater than 1. Thus, we propose a dynamic estimator in which the denominator is calculated by defining a donor population (P_D_) for each site independently. For each site, P_D_ is the population (either P_2_ or P_3_) that has the higher frequency of the derived allele, thus maximizing the denominator and eliminating *f* estimates greater than 1:

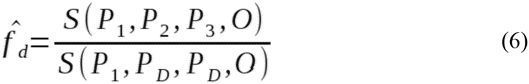

### Assessing the ability of *D* and ***f*** estimators to quantify introgression in small sequence windows

To assess how reliably Patterson’s *D* statistic, and other estimators of *f* are able to quantify the actual rate of introgression, we simulated sequence datasets with differing rates of introgression using ms (Hudson 2002). For each dataset, we simulated 100 sequence windows for 8 haplotypes each from four populations with the relationship (((P_1_,P_2_),P_3_),O). The split times *t_12_* and *t_23_* (as on Figure 1A) were set to 1 × 4*N* generations and 2 × 4*N* generations ago, respectively, and the root was set to 3 × 4*N* generations ago. An instantaneous, unidirectional admixture event, either from P_3_ to P_2_ or from P_2_ to P_3_, was simulated at a time *t_GF_* with a value *f*, which determines the probability that each haplotype is shared. We tested two different values for *t_GF_*: 0.1 and 0.5 × 4*N* generations ago. For each direction of gene flow and each *t_GF_*, 11 simulated datasets were produced, with *f* values ranging from 0 (no gene flow) to 1 (all haplotypes are shared). Finally, the entire set of simulations was repeated with three different window sizes: 1, 5 and 10 kb, and with three different recombination rates: 0.001, 0.01 and 0.1, in units of 4Nr, the population recombination rate. DNA sequences were generated from the simulated trees using Seq-Gen (Rambaut & Grass 1997), with the HKY substitution model and a branch scaling factor of 0.01. Simulations were run using the provided script compare_f_estimators.r, which generates the ms and Seq-Gen commands automatically. An example set of commands to simulate a single 5kb sequence using the split times mentioned above, with gene flow from P_3_ to P_2_ at t_GF_ = 0.1 and *f* = 0.2, and with a recombination rate parameter of 0.01 would be:

~~~
ms 32 1 −I 4 8 8 8 8 −ej 1 2 1 −ej 2 3 1 −ej 3 4 1 −es 0.1 2 0.8 −ej 0.1 5 3 −r
50 5000 −T | tail −n + 4 | grep −v // > treefile
partitions=($(wc −l treefile))
seq−gen −mHKY −l 5000 −s 0.01 −p $partitions < treefile > seqfile
~~~

We then compared the mean and standard error for *D* (Equation 1) and the three *f* estimators (Equations 4, 5 and 6), calculated for all 100 windows in each dataset.

### Analysis of *Heliconius* whole genome sequence data

To investigate how the *D* and 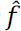 statistics are affected by underlying diversity in a given window, we re-analyzed whole genome data from Martin et al. (2013). For ABBA-BABA analyses, populations were defined as follows: P_1_ = *Heliconius melpomene aglaope* (4 diploid samples), P_2_ = *Heliconius melpomene amaryllis* (4), P_3_ = *Heliconius timareta thelxinoe* (4), O = *Heliconius hecale* (1), *Heliconius ethilla* (1), *Heliconius pardalinus sergestus* (1), and *Heliconius pardalinus* ssp. nov. (1). Patterson’s *D* (Equation 1) and the three *f* estimators (Equations 4,5,6) were calculated, along with nucleotide diversity (π) and absolute divergence (*d_XY_*), for non-overlapping 5 kb windows across the genome. Both π and *d_XY_* were calculated as the mean number of differences between each pair of individuals, sampled either from the same population (π), or from separate populations (*d_XY_*). Sites with missing data were excluded in a pairwise manner, and each pair of individuals contributed equally to the mean. Windows were restricted to single scaffolds and windows for which fewer than 3000 sites had genotype calls for at least half of the individuals were discarded. To calculate *D* and the *f* estimators only bi-allelic sites were considered. The ancestral state was inferred using the outgroup taxa, except when the four outgroup taxa were not fixed for the same allele, in which case the most common allele overall was taken as ancestral. The *HmB* locus was defined as positions 300 000 to 450 000 on scaffold HE670865 and the *HmYb* locus as positions 650 000 to 900 000 on scaffold HE667780 of version 1.1 of the *H. m. melpomene* genome sequence. We also analyzed windows from each of the 21 chromosomes of the *H. m. melpomene* genome sequence separately. Scaffolds were assigned to chromosomes according to the *Heliconius* Genome Consortium (2012), and incorporating the improved assignment of Z-linked scaffolds by Martin et al. (2013) (details available in Dryad repositories http://dx.doi.org/10.5061/dryad.m27qq and http://dx.doi.org/10.5061/dryad.dk712). This analysis was performed using egglib_sliding_windows.py, and figures were generated using Figures_3_S3_S4.R.

### Assessing a test to distinguish introgression from shared ancestral variation based on absolute divergence

Smith and Kronforst (2013) proposed a simple test to distinguish between the hypotheses of pre- and post-speciation shared ancestry based on absolute divergence. To assess this method on data of known history, we generated a large range of sequence datasets using ms (Hudson 2002) and Seq-Gen (Rambaut & Grass 1997). For the simplest (‘null’) model 10 000 5kb sequence windows were simulated for 8 haplotypes each from three populations and an outgroup, with the relationship (((P_1_,P_2_),P_3_),O), without gene flow or population structure. To approximate a scenario in which a subset of the genome has a distinct phylogenetic history, either due to gene flow or genomically-localized ancestral population structure, we used a combined model approach. This entailed combining 9000 5kb windows from the null model (90% “Background” windows), with 1000 5kb windows simulated under with the topology ((P_1_,(P_2_,P_3_)),O), consistent with shared ancestry between P_2_ and P_3_ (10% “Alternate” windows). By altering the split times, three distinct scenarios were emulated: Gene flow from P_2_ to P_3_, gene flow from P_3_ to P_2_, and ancestral structure (Figure 4A-D). Using entirely distinct topologies in this way is equivalent to making the probability of gene flow (or structure) equal to one in the 1000 Alternate windows. While this approach of partitioning each dataset into two somewhat arbitrarily-sized subsets with evolutionary histories at two extremes is biologically unlikely, it provided a simple and powerful framework in which to evaluate Smith and Kronforst’s approach, with clear expectations. Model combination datasets were generated using run_model_combinations.py and shared_ancestry_simulator.R, which generates the ms and Seq-Gen commands automatically, in a similar form to those given above. For example if *t_12_* = 1, *t_23_* = 2, 4Nr = 0.01 and gene flow from P_3_ to P_2_ at *t_GF_* = 0.2, the ms calls for Background and Alternate models, respectively, would be:

~~~
ms 32 1 −I 4 8 8 8 8 −ej 1 2 1 −ej 2 3 1 −ej 3 4 1 −r 50 5000 −T
ms 32 1 −I 4 8 8 8 8 −ej 0.2 2 3 −ej 2 3 1 −ej 3 4 1 −r 50 5000 −T
~~~

We calculated Patterson’s *D* (Equation 1) and the three *f* estimators (Equations 4,5,6) for all windows, and identified the top 1000 ‘outliers’ (10%) with the most extreme values. For *D*, only positive values were included as outliers, as negative values indicate an excess of BABAs, consistent with introgression between P_1_ and P_3_. Similarly, for *f* estimators, only windows with *D* ≥ 0 were considered, as these values only give meaningful quantification of introgression when there is an excess of ABBAs. To compare P_2_-P_3_ divergence between the Background and Alternate windows, or between outlier and non-outlier windows, we calculated *d_XY_* for each window as described above, for each pair of populations. Average *d_XY_* was compared between subsets of windows using a Wilcoxon rank-sum test, as values tended to be non-normally distributed (confirmed with Bonferroni-corrected Shapiro-Wilk tests).

These tests were repeated over a large range of split times. In all cases the root was set to 3.0 × 4*N* generations ago, and the other splits ranged from 0.2 to 2.0. Times of gene flow and structure also varied same scale. In total this gave 45 null models and 120 models each for the two gene flow scenarios and ancestral structure (405 overall). The analyzed models therefore covered a vast range of biologically relevant scales. In all cases, the Seq-Gen branch scaling factor was set to 0.01. Full parameters for all models are provided in Table S1. Finally, to examine the effects of recombination rate, the entire simulation study was repeated using population recombination rate (4Nr) values of 0.01 and 0.001. Summary statistics for all models were compiled using generate_summary_statistics.R.

### Software

Code and data for this manuscript will be made available as a Data Dryad repository on acceptance. Most files, with instructions for running scripts to generate the results, are currently available on GitHub at https://github.com/johnomics/Martin_Davey_Jiggins_evaluating_introgression_statistics. Large data sets can be made available on request and will be made publicly available on Data Dryad on acceptance. This work was made possible by the free, open source software packages EggLib (De Mita and Siol 2012), phyclust (Chen 2011), R (R Core Team 2013), ggplot2 (Wickham 2009), plyr (Wickham 2011), reshape (Wickham 2007) and Inkscape (http://www.inkscape.org).

## ACKNOWLEDGEMENTS

We are grateful to Milan Malinsky for helpful and detailed discussions about this work. We also thank Jim Mallet, Marcus Kronforst and two anonymous reviewers for their comments on the manuscript. Richard Merrill and Richard Wallbank contributed to initial discussions that inspired the work. This work was supported by the Leverhulme Trust (F/09364/E to C.D.J.) and the Herchel Smith Fund (Postdoctoral Fellowship to J.W.D.).

